# TranslAGE: A Comprehensive Platform for Systematic Validation of Epigenetic Aging Biomarkers

**DOI:** 10.1101/2025.10.16.682960

**Authors:** Daniel S. Borrus, Raghav Sehgal, Jenel F. Armstrong, John Gonzalez, Grace Zou, Jessica Kasamoto, Yaroslav Markov, Jessica Lasky-Su, Albert Higgins-Chen

## Abstract

Epigenetic clocks are powerful biomarkers of biological aging, however, their performance varies across studies and contexts. Current limitations include siloed datasets, inconsistent validation methods, and the absence of a standardized framework for systematic comparison. Here, we introduce TranslAGE: a publicly available online resource that addresses this gap by harmonizing 179 human blood DNA methylation datasets and precalculating a suite of 41 epigenetic biomarker scores for each of the >42,000 total samples. Users can explore these data through interactive dashboards that evaluate four fundamental performance domains: Stability, Treatment response, Associations, and Risk, collectively forming the STAR framework. Stability quantifies robustness to multiple types of technical and biological noise. Treatment response measures biomarker sensitivity to aging interventions and environmental exposures. Associations capture cross-sectional relationships with age, demographics, disease, and other phenotypes, and Risk assesses predictive power for future functional decline, morbidity and mortality. The STAR framework unifies these test metrics into a single composite scoring system that enables researchers to identify, benchmark, and validate biomarkers best suited to their scientific or clinical applications. TranslAGE will be continually updated, with rapid scaling by adding datasets, biomarkers, or analyses. By providing harmonized datasets, precomputed biomarker scores, and interactive data tools, TranslAGE establishes the first standardized, reproducible framework for benchmarking epigenetic aging biomarkers across populations, and accelerates the translation toward clinical use.

## Introduction

Epigenetic-based biomarkers represent an exciting new tool in the quest for longer, healthier lives (Duan et al. 2022; Horvath and Raj 2018). By estimating biological age, they are strong predictors of aging outcomes, such as cardiovascular disease, cancer, and all-cause mortality (Fransquet et al. 2019; Simpson and Chandra 2021; Oblak et al. 2021). Integrating both hereditary and environmental influences, these measurements provide a potential framework for monitoring and altering aging trajectories (A. Li, Koch, and Ideker 2022; Quach et al. 2017; Salameh, Bejaoui, and El Hajj 2020). With over a million detectable methylation sites in modern arrays, the majority of which change with age (Seale et al. 2024; Jacques et al. 2025), new clocks are developed rapidly. There are good reasons for having many clocks: initially clocks advanced from predicting chronological age (Horvath 2013; Hannum et al. 2013) to predicting mortality risk or pace of aging (Lu et al. 2019; Levine et al. 2018; Belsky et al. 2022). Later clocks aimed to disentangle specific aspects of aging, such as aging in different physiological systems, damaging versus adaptive changes, or specific outcomes like frailty (Ying, Liu, et al. 2024; Sehgal, Markov, et al. 2025; X. Li et al. 2022). Additional clocks have been developed for specific tissues, populations, age ranges, or applications (Galkin et al. 2023; Bell et al. 2019). The number of algorithms will continue to expand, as thousands of methylation-based algorithms for different aspects of aging biology have recently been reported (Ying, Tyshkovskiy, et al. 2024; Carreras-Gallo et al. 2024). This raises a deceptively simple question for researchers: “What clock should I use?”

Epigenetic clocks have been well-studied as a result of widespread DNAm data and the relative ease of training and applying new algorithms. The latest meta-analysis of DNAm aging biomarker associations included 299 publications and 1,050 phenotypes (Chervova et al. 2024). DNAm has been assayed in a variety of contexts, including many cohorts with long-term aging outcomes and studies of geroscience interventions (Moqri et al. 2023; Moqri, Herzog, et al. 2024). Importantly, the frequent use of Illumina BeadChip methylation arrays measuring a common set of CpGs means that methylation data can typically be compared across studies. Most DNAm biological age algorithms are compatible across 450K, EPICv1, and EPICv2 arrays. Thus, with DNAm it is possible to simultaneously compare a DNAm biomarker’s prediction of aging outcomes in many cohorts and its response to a variety of interventions, although this promise has not yet been realized.

A major challenge is that the large number of both clocks and DNAm datasets leads to a highly fragmented landscape of epigenetic biomarkers that makes it very difficult for researchers to select a biomarker for study. Many studies examine only one or a handful of biomarkers, while those that compare many will calculate inconsistent sets of clocks and introduce multiple testing issues. A systematic review reveals highly uneven deployment of clocks in different studies examining their phenotypic associations (Chervova et al. 2024), and we previously summarized a similar trend in intervention studies (Sehgal et al. 2024). Even when studies attempt to comprehensively characterize many clocks, this work necessarily becomes out of date rapidly because of the constant development of new clocks. When different studies disagree on a result, it is unclear whether this is because of differences in clock selection, true biological differences in the study populations, or variations in analytical approach. Furthermore, much of the epigenetic clock literature involves clocks being applied to one dataset from the massive amounts of publicly available methylation data from public repositories, which risks publication bias if one clock happens to show a significant phenotypic association in one dataset by chance. Compounding the issue of numerous clocks is the fact that there are numerous potential metrics to evaluate a given biomarker, including responsiveness to many interventions, prediction of diverse aging outcomes, and robustness to various technical and biological factors that may modify clock performance. Many studies examine one property of biomarkers at a time, meaning each discovery exists in isolation even if multiple clocks are compared. There are also many possible applications of clocks (forensics, epidemiology, basic geroscience, comparative biology, clinical trials, longitudinal tracking, etc.), and it may be the case that different clocks are most appropriate for different use cases. Taken together, researchers are faced with myriad choices for clocks and must wade through a highly confusing literature with numerous clocks, various biological and technical considerations, inconsistent application of clocks, and out-of-date information, making it nearly impossible to select the best ones when planning their own study.

Cross-study comparison is essential to reinforce findings, demonstrate robustness across study design, and build field-wide consensus on the specific capabilities of different aging biomarkers (Moqri, Herzog, et al. 2024). Several packages now allow calculation of a large number of clocks in a harmonized manner, such as methylCIPHER, Biolearn, Pyaging, methScore, and EpigeneticAgePipeline (Thrush et al. 2022; Moqri, Ying, et al. 2024; de Lima Camillo 2023; Xu et al. 2024; Rayevskiy et al. 2024). There have also been efforts to harmonize methylation datasets (S. Li et al. 2025; Maden et al. 2023; Xiong et al. 2020). A few efforts have been made to systematically benchmark clocks (Chervova et al. 2024; Ying et al. 2023; Kriukov et al. 2024), but the metrics are largely restricted to associations, without benchmarking of reliability or intervention response. There is no resource where all of these clock algorithms and datasets have already been analyzed to provide the comprehensive insights about each clock’s performance needed to select a clock.

These gaps underscore the need for a unified benchmarking system that tests robustness, responsiveness, and predictive validity across multiple contexts and many studies. (Biomarkers of Aging Consortium et al. 2024). Here, we present TranslAGE which was developed to provide exactly this. It is a platform for harmonizing methylation datasets and precalculating a standardized suite of epigenetic clock scores, enabling systematic comparison across a range of metrics. At launch, TranslAGE measures and compares biomarkers across four core domains that together form each clock’s STAR score: Stability to biological and technical noise, Treatment response to aging interventions or adverse events, Associations with aging phenotypes and diseases, and Risk predictions for future disease or mortality. This framework allows for robust cross-validation of epigenetic biomarkers, bridging the gaps in the current landscape. Both domain-specific results and summarized STAR scores, along with all harmonized datasets and precalculated clock outputs, are accessible through TranslAGE’s interactive online dashboards.

## Results

### TranslAGE Resource Overview

At the time of submission, TranslAGE integrates 179 methylation datasets, a number that continues to expand as new studies are harmonized. Across all datasets, TranslAGE encompasses >34,000 unique individuals and >42,000 methylation profiles, spanning both Illumina 450k and EPIC arrays. Of these, 129 datasets are longitudinal and 51 are cross-sectional. Supplementary Table 1 contains the complete list of datasets at time of this submission.

For each dataset, we precalculated 41 epigenetic clocks, spanning first-generation chronological age predictors (Hannum et al. 2013; Horvath 2013), second generation mortality clocks (Lu et al. 2019; Belsky et al. 2022, 2020; Levine et al. 2018), reliable clocks (Higgins-Chen et al. 2022), and system-specific ages (Sehgal, Markov, et al. 2025). We calculate nearly all publicly available epigenetic clocks, and are capable of rapidly incorporating new algorithms or those provided by collaborators. The breadth of epigenetic-based algorithms in TranslAGE enables systematic evaluation of aging biomarkers at scale not previously possible.

Figure 1 provides an overview of the initial processing steps of the TranslAGE data harmonization framework. We compiled a large collection of methylation datasets from public sources, including NCBI GEO (Clough et al. 2024) and EMBL (Athar et al. 2019), as well as private datasets shared by collaborators. To maximize initial utility for clinical trial planning, we required eligible datasets to meet three inclusion criteria: 1) human origin, 2) generation from a methylation array platform, and 3) derived from whole-blood samples which facilitates longitudinal monitoring.

**Figure 1:**
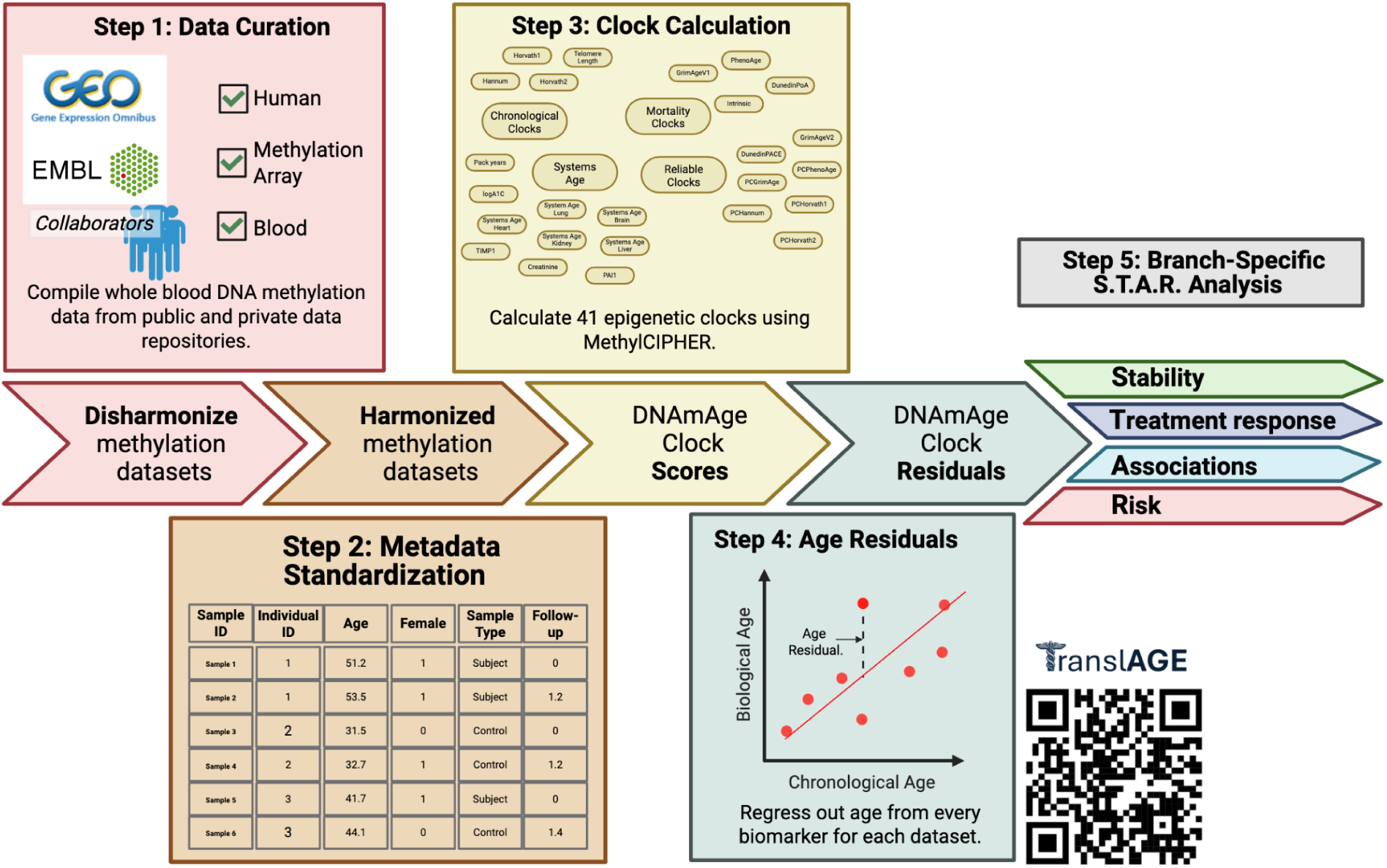
Methodology Overview. The TranslAGE work flow begins with data curation, where whole-blood methylation datasets are gathered from public and private sources (e.g., GEO, EMBL, TruDiagnostic). Next, a metadata standardization step harmonizes sample identifiers, subject IDs, chronological age, sex, group labels and follow-up times across datasets (illustrated by a standardized table). The clock calculation step uses the methylCIPHER package to compute 41 epigenetic clocks, including first-generation clocks, second-generation clocks and various multi-omic signatures. These clock scores are then residualized for age and sex to produce age residuals (shown as the vertical difference between biological age and chronological age on a scatter plot). Finally, the pipeline branches into the four STAR analyses (Stability, Treatment Response, Associations, and Risk).

To enable cross-study comparability, we standardized all datasets to a shared metadata schema. Each dataset was reformatted to include a uniform set of variables. These curated columns include Sample ID, Subject ID, Sex, Age, Group, and Time of Sample From Baseline. This harmonization allows any dataset within TranslAGE to pass through identical processing and analysis workflows.

Each dataset was then classified according to its suitability for one or more of the four TranslAGE analysis domains. Datasets containing baseline or future clinical phenotypes and disease outcomes were routed to the Association and Risk domains, respectively. Datasets containing technical replicates or repeated samples within a 24 hour window (biological replicates) were assigned to the Stability domain. Some stability datasets included events that are inevitable parts of daily life (meals, mild stress, exposure to pollen) in between time points. Longitudinal dataset with repeated measures from the same individual were assigned to the Treatment response domain. Most Treatment response datasets contained a major intervention (e.g. new diet and exercise regimen, new metformin treatment) or major event (new chemotherapy, new-onset dementia). However, longitudinal datasets where no clear event or health change occurs were also assigned to Treatment response to serve as controls.

For every eligible dataset, we computed a standardized suite of 41 epigenetic clocks. This pipeline is an extension of the core framework of the methylCIPHER R package (available at https://github.com/HigginsChenLab/methylCIPHER), our package for calculating epigenetic clocks. Supplementary Table 2 contains the list of clocks present in TranslAGE at time of this submission. Each biomarker score (DNAmAge) was subsequently residualized for chronological age and sex, enabling comparison across individuals.

Following residualization, the resulting DNAmAge residuals were dispatched to their designated analysis pipelines (Stability, Treatment response, Associations, and Risk) depending on the dataset classification. A complete list of harmonized datasets, their metadata, and assigned analysis domains is available through the TranslAGE Front Page Dashboard (translage.io/frontpage). Additionally, full metadata, harmonized datasets and precalculated biomarkers are accessible for download (*pending publication in a peer-reviewed Journal, limited to datasets that were originally public*). Figure 2 illustrates the potential use case of a clinical researcher planning a new lifestyle intervention trial with the TranslAGE Front Page Dashboard.

**Figure 2:**
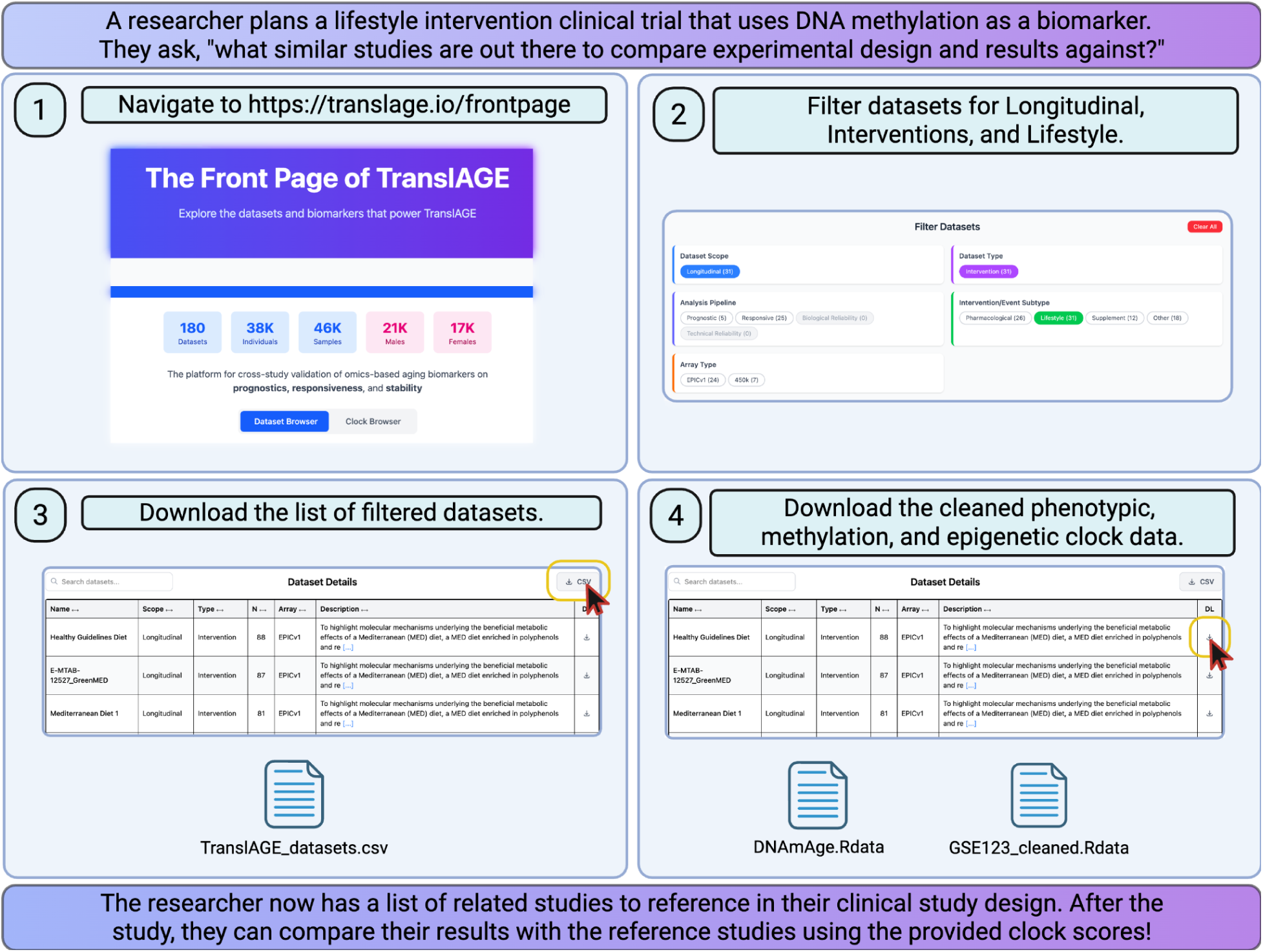
Front Page Use Case. A tutorial for using the Front Page Dashboard of TranslAGE, as a way to identify and filter potential methylation datasets. Screenshots are directly from the TranslAGE website. Yellow boxes and red-lined cursors are used to highlight important elements of the interactive page.

### Stability to Confounders

Quantifying biomarker stability is essential for determining which measures remain robust to technical and biological sources of noise. This is particularly an issue if the magnitude of noise outweighs the effect of interventions under study, which can dramatically increase the required sample size to detect intervention effects(Higgins-Chen et al. 2022; Sehgal, Borrus, et al. 2025). To evaluate reliability, we constructed a stability pipeline encompassing both technical and biological replicate datasets.

Technical replicates were defined as samples obtained from the same individual and the same blood draw, capturing variability introduced by array processing or batch effects. Biological replicates were defined as multiple samples collected from the same individual within a short time window, where natural events of daily life (mild stress, circadian rhythm, dietary intake, pollen exposure) could influence methylation profiles.

In total, five datasets contained technical replicates and six datasets contained biological replicates. For each clock within each dataset, we computed the intraclass correlation coefficient (ICC) to quantify the proportion of total variance attributable to between-individual, rather than within-individual, differences. To pool results across datasets, ICCs were transformed using Fisher’s z-transformation and combined through random-effects meta-analysis (see *Methods*).

We calculated pooled technical and biological ICCs for each of the 41 epigenetic clocks, providing dual stability metrics for each biomarker. These results form the foundation of the Stability Dashboard (translage.io/stable), which enables interactive exploration and ranking of biomarkers by technical and biological reliability. Figure 3 illustrates the stability analysis workflow, and the following use cases demonstrate how TranslAGE identifies biomarkers that are robust, or sensitive, to specific sources of experimental noise.

**Figure 3:**
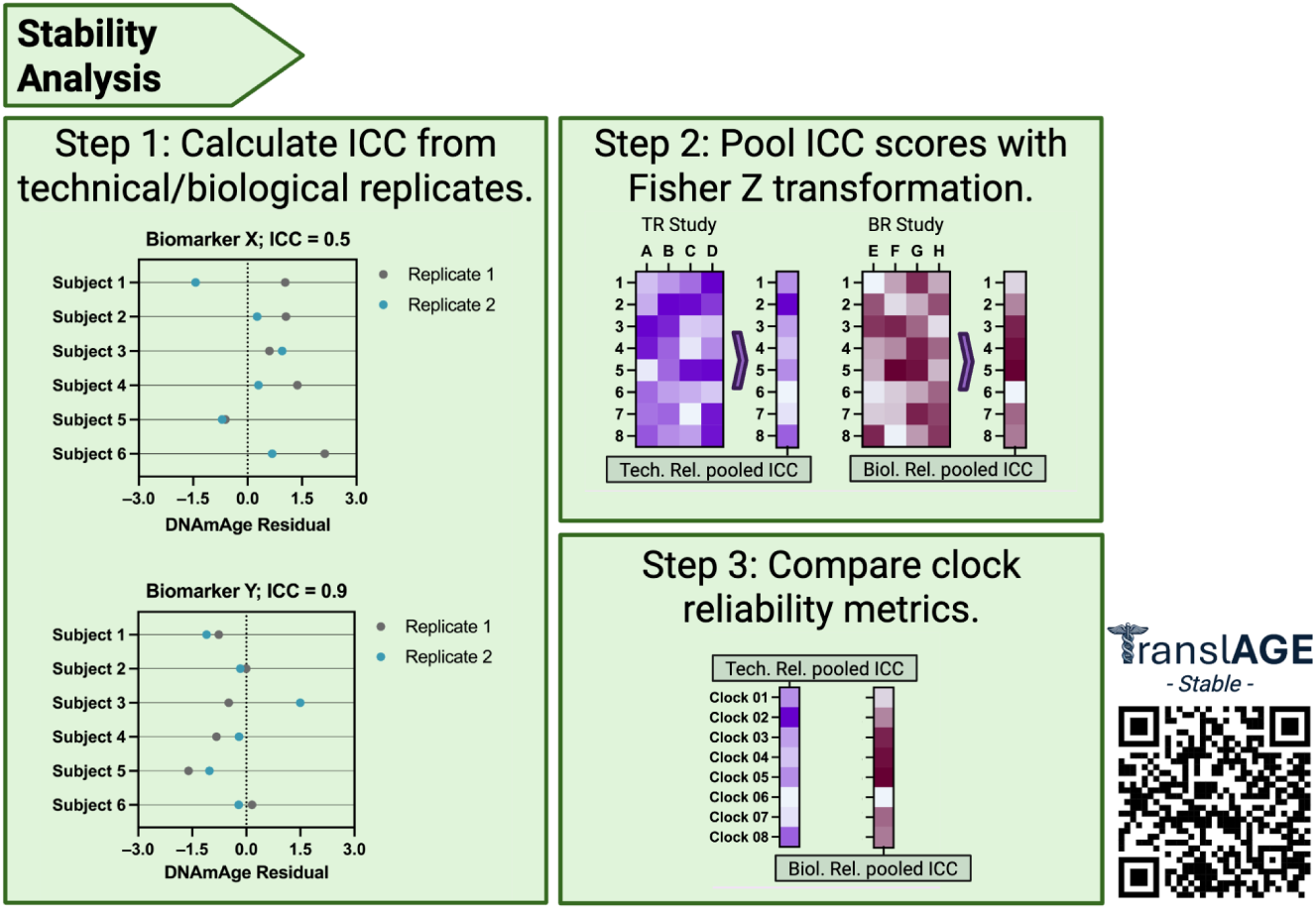
Stability Analysis of Epigenetic Clocks. This figure shows how the pipeline assesses the technical and biological reliability of clocks. **Step 1** calculates intraclass correlation coefficients (ICCs) using technical or biological replicate samples; scatter plots show biomarker X (ICC = 0.5) and biomarker Y (ICC = 0.9), with replicate 1 and 2 values for each subject plotted on the same axis to illustrate lower and higher repeatability. **Step 2** pools ICCs across studies using Fisher-Z transformation; heatmaps depict technical-replicate (TR) and biological-replicate (BR) studies, where each cell represents a clock’s ICC in a given study. **Step 3** compares pooled technical and biological reliability metrics across clocks; a bar-like heatmap shows how specific clocks rank in technical versus biological ICCs, enabling identification of robust biomarkers. The QR code directs to translage.io/stable.

**Figure 4:**
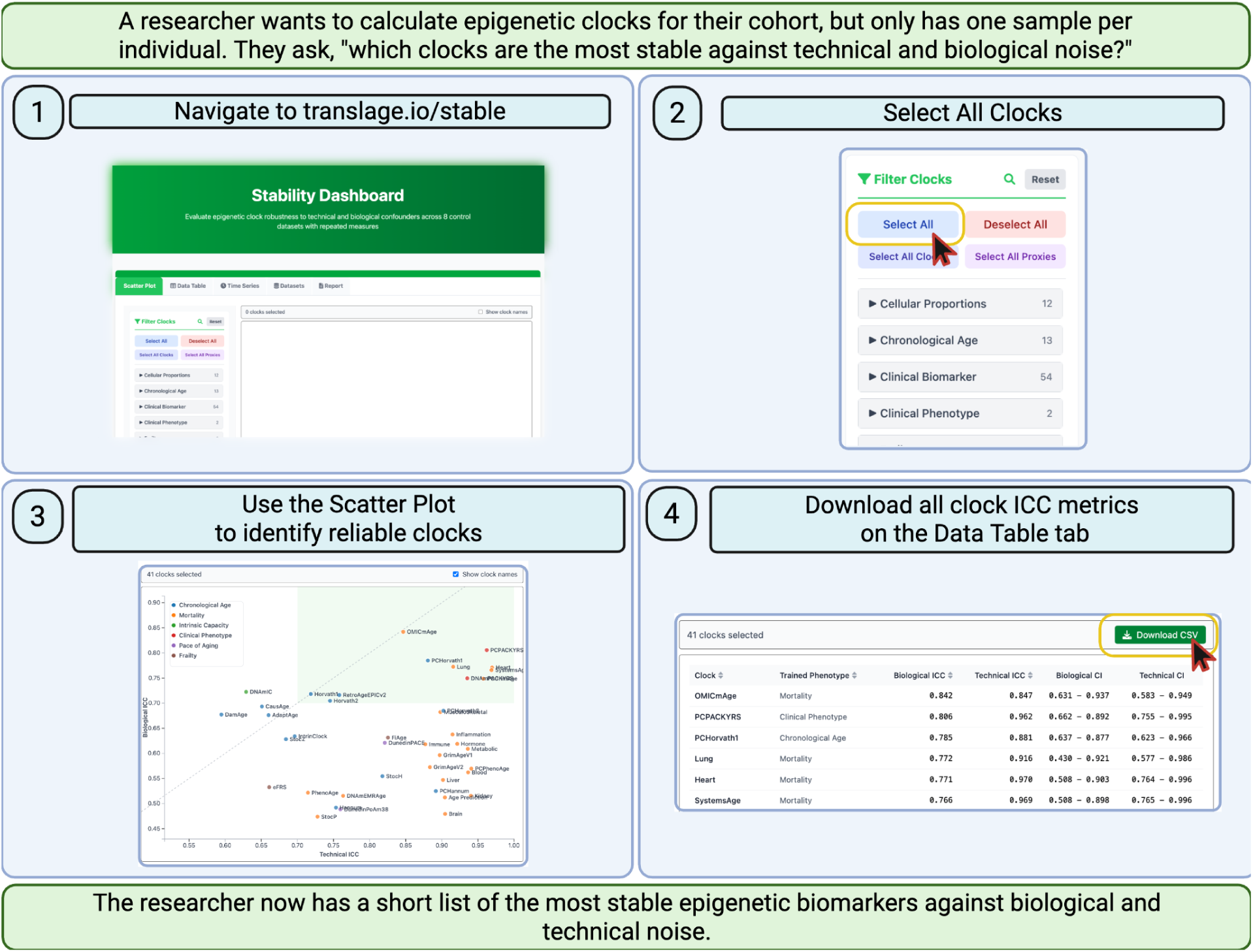
Stability Use Case. **A.** A tutorial on using the Stability Dashboard to investigate clock reliability, and filter for the most stable clocks. Screenshots are directly from the TranslAGE website. Yellow boxes and red-lined cursors are used to highlight important elements of the interactive page.

**Figure 5:**
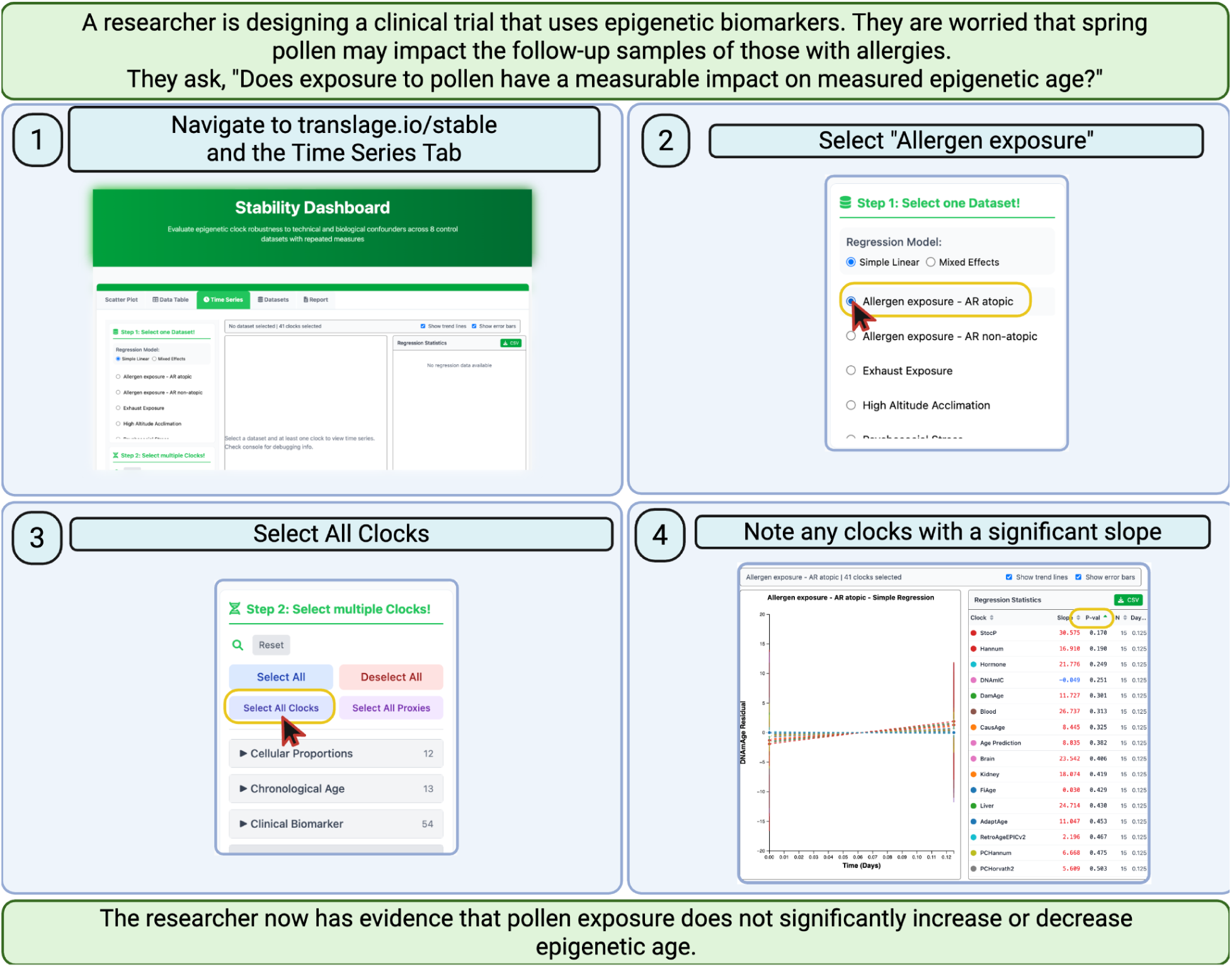
Stability Use Case. **B.** A tutorial on using the TranslAGE Stability to investigate biological confounders and their measurable impact on epigenetic age. Screenshots are directly from the TranslAGE website. Yellow boxes and red-lined cursors are used to highlight important elements of the interactive page.

### Biomarker Treatment Response to Interventions and Aging Events

Longitudinal datasets containing repeated samples from the same individuals enabled investigation of how each biomarker responds to beneficial or adverse aging-related events. Among the 129 longitudinal datasets in TranslAGE, 101 of them passed quality control for the Treatment response pipeline (see *Methods*). Of these 101 datasets, 74 contained paired methylation information before and after a putative aging intervention, such as a Mediterranean diet, smoking cessation, or initiation of metformin. This list includes datasets from our prior harmonized analysis of 51 different putative longevity interventions and their effects on DNAm aging biomarkers (Sehgal et al. 2024). We further subdivided the aging interventions into Lifestyle (25 datasets), Supplement (11 datasets), Pharmacological (22 datasets), and Other (16 datasets). Additionally, 19 datasets captured a potentially adverse exposure, such as radiotherapy, SARS-CoV-2 infection, or air pollution. 8 datasets contained longitudinal methylation data from a control study, in which no experimentally applied conditions were present.

To standardize comparison across studies, we conducted paired t-tests comparing each subject’s baseline DNAmAge residuals to those at follow-up. For every dataset-biomarker combination, we extracted the mean effect size and the corresponding test statistics (including p-values). Negative effect sizes indicate a decrease in biological age (beneficial effect); a positive effect size indicates an increase in biological age (deleterious effect).

Because each clock differs in magnitude of scale, all effect sizes were standardized to the population-level DNAmAge standard deviation from a large EPICv1 dataset. Under this scaling, an effect size of 1.0 represents a one-standard deviation increase in DNAmAge. Additional details of the analysis pipeline are provided in the *Methods* section.

All scaled effect sizes and associated p-values are available through the Treatment Response Dashboard (translage.io/responsive), which allows users to explore how different interventions or exposures modulate specific epigenetic clocks. Figure 6 illustrates the analytic flow of the Treatment response pipeline. Figure 7 highlights a use case of how TranslAGE can help a researcher identify responsive clocks and the interventions that target them.

**Figure 6:**
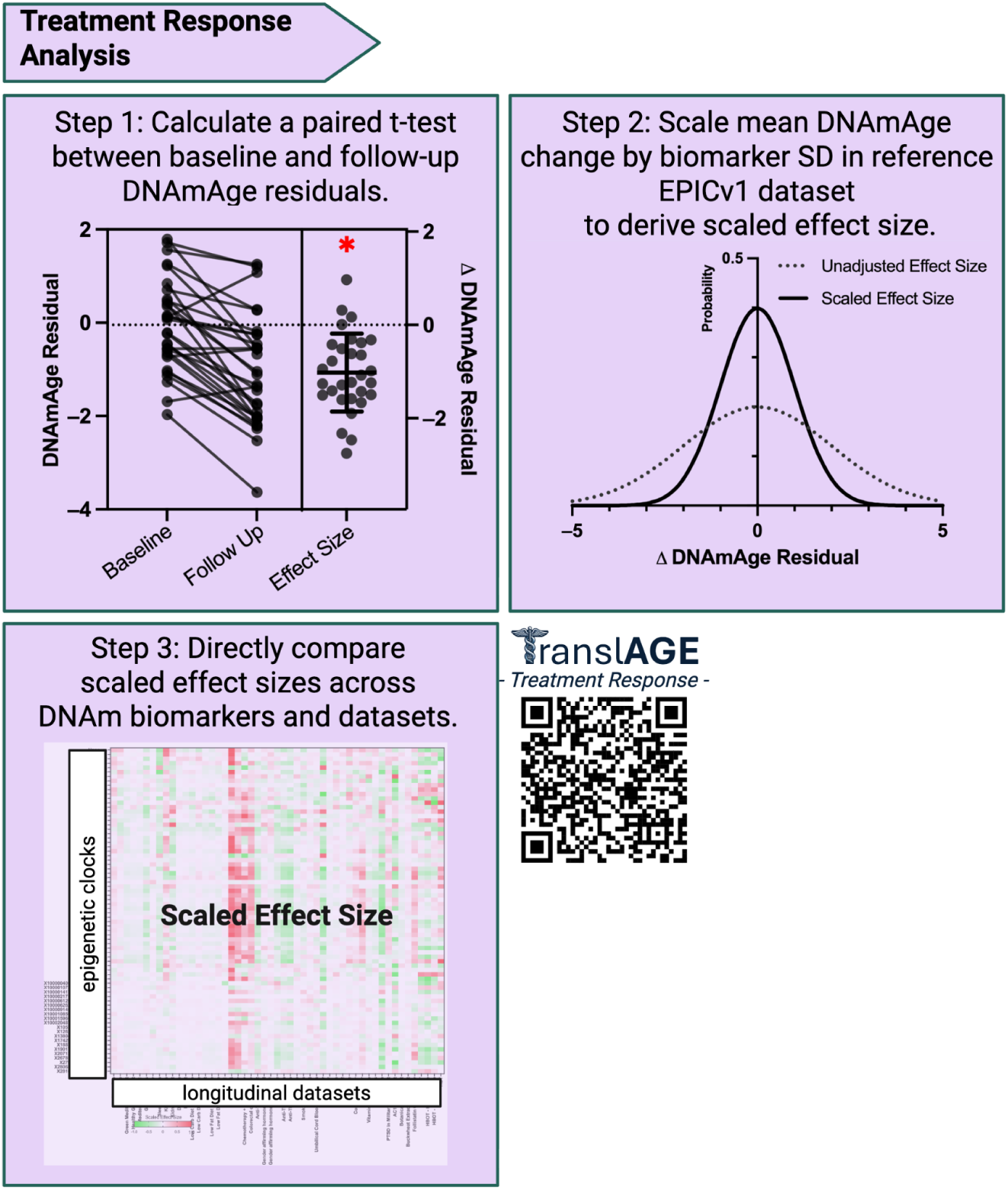
Methods for Treatment Response Analysis. This figure outlines how TranslAGE assesses whether biomarkers respond to interventions or adverse events. **Step 1** performs paired t-tests on DNAmAge residuals before and after an intervention; the schematic shows baseline and follow-up residuals for multiple subjects connected by lines, with a distribution of the resulting effect sizes. **Step 2** scales the mean change in DNAmAge by the biomarker’s age-adjusted standard deviation in a large EPICv1 dataset to derive a standardized effect size; the schematic contrasts unadjusted and scaled effect size distributions. (SD = Standard Deviation). **Step 3** compares scaled effect sizes across all biomarkers and datasets; a heatmap displays effect sizes from longitudinal datasets, illustrating the large-scale comparison of biomarker responsiveness. The QR code directs to translage.io/responsive.

**Figure 7:**
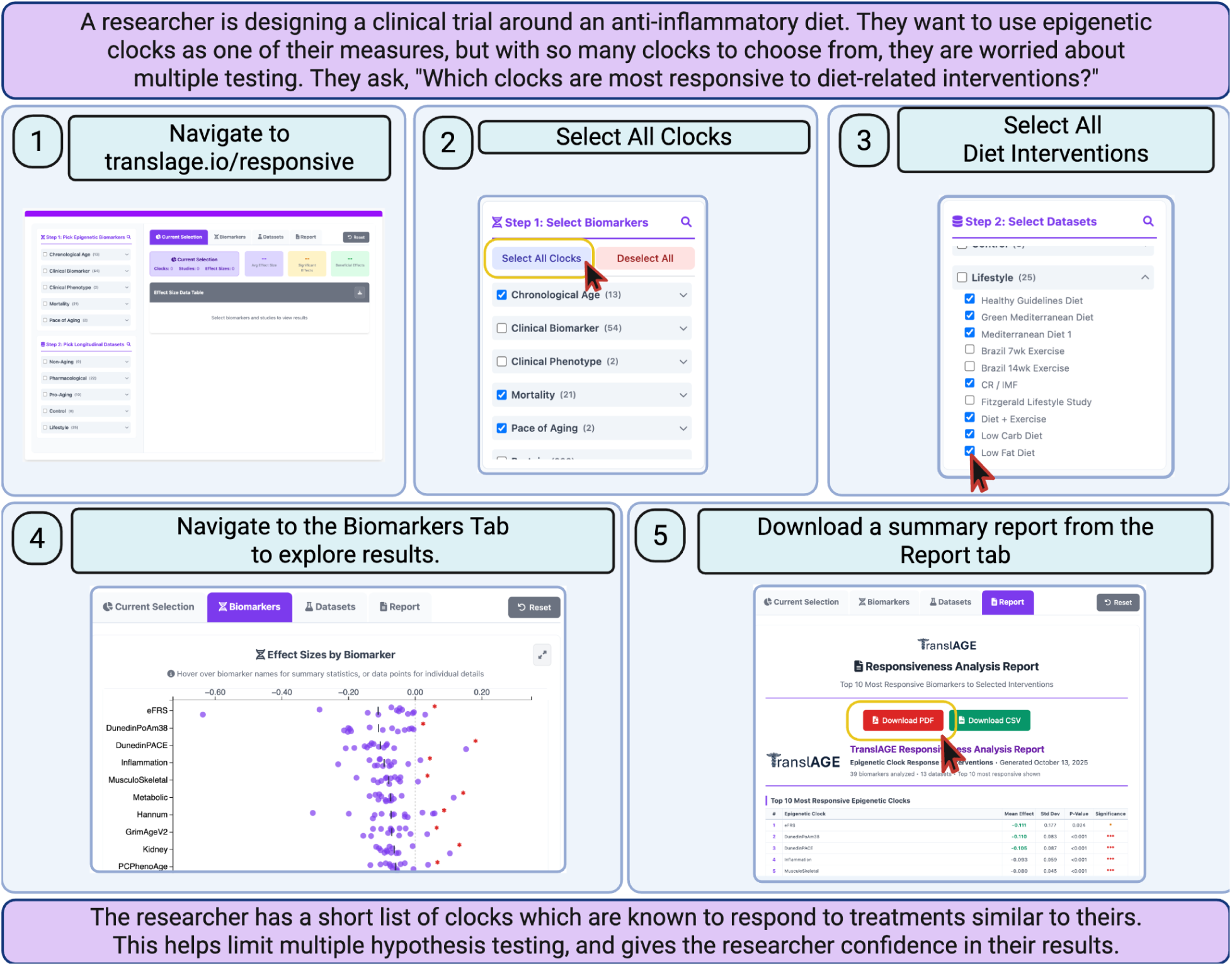
Treatment Response Use Case. **A:** tutorial showcasing how to use the TranslAGE Treatment Response Dashboard to determine the most responsive clocks to diet interventions. Screenshots are directly from the TranslAGE website. Yellow boxes and red-lined cursors are used to highlight important elements of the interactive page.

### Association with Aging-Related Phenotypes and Disease

During dataset harmonization, phenotypic variables related to disease and aging were identified and reformatted into a standardized structure. Among the 179 datasets in TranslAGE, 18 of them contained baseline health information, including concurrent disease diagnosis and clinical biomarker measurements. 16 of these datasets shared at least one overlapping phenotype, enabling cross-study meta-analysis. In total, 120 dataset-phenotype pairs were identified across the 18 datasets, with 22 baseline phenotypes having association information from more than 1 dataset. Supplementary Table 3 contains details on phenotype definitions and categorizations.

For each dataset, we performed association analysis between DNAmAge residuals and available baseline phenotypic variables. Model choice was determined by phenotype category. Binary outcomes, such as disease diagnosis, were analyzed with logistic regression. Continuous phenotypes, such as BMI or blood pressure, were analyzed with linear regression. All models were adjusted for chronological age and sex.

To aggregate findings across datasets measuring the same phenotype, we conducted random-effects meta-analysis to compute pooled effect sizes for each clock-phenotype pair. For binary disease statuses, we additionally combined group means and standard deviations for case versus control groups using meta-analytic formulas (see *Methods*).

After meta-analysis, TranslAGE included 90 unique phenotypes. Figure 8 summarizes the workflow of this analysis. These aggregated results can be explored interactively through the Associations Dashboard at translage.io/associations, which allows users to query baseline associations by phenotype or biomarker. Figure 9 depicts a possible use case for the Associations Dashboard.

**Figure 8:**
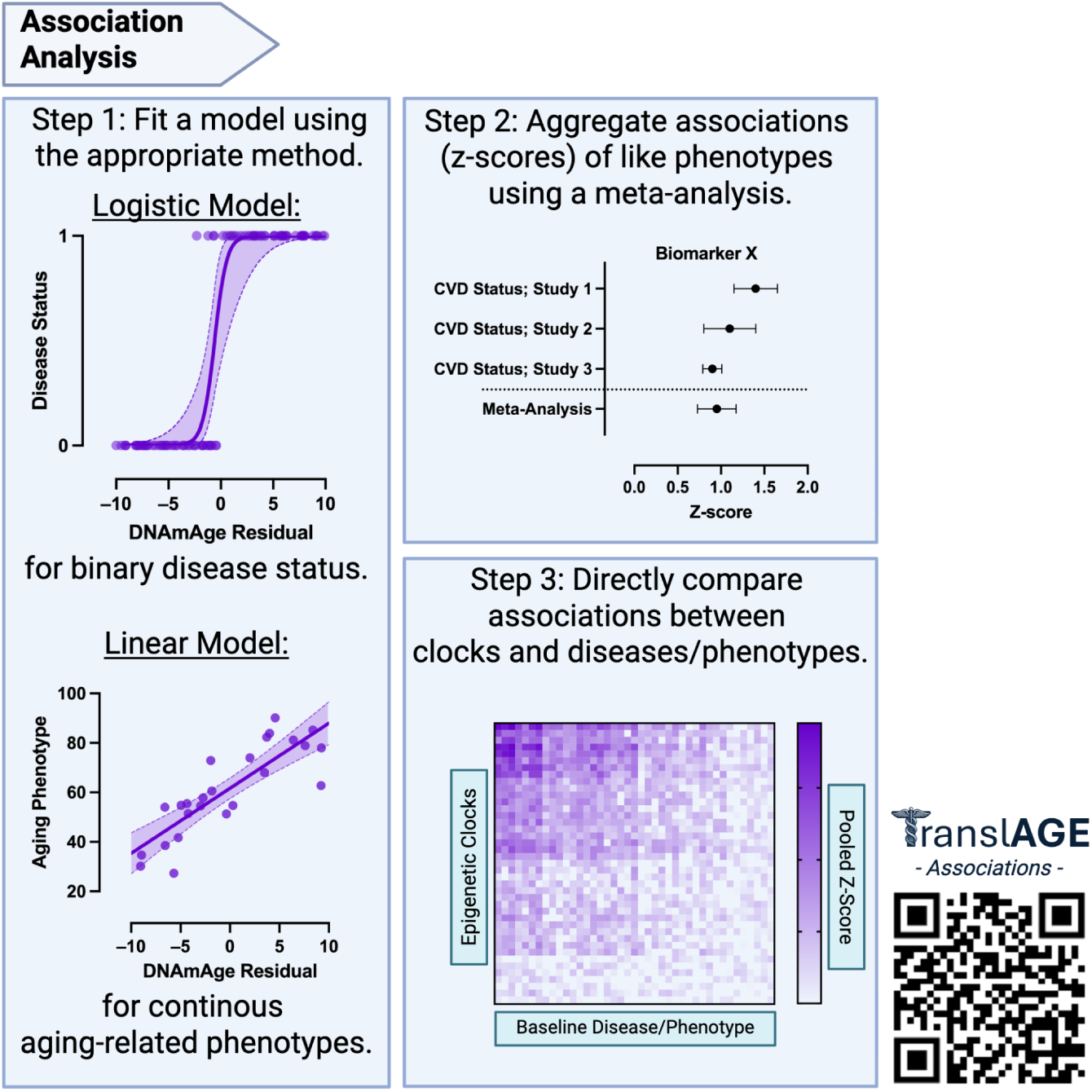
Associations Analysis Pipeline. This figure illustrates how TranslAGE evaluates biomarkers associations with baseline disease or aging phenotypes. **Step 1** fits a regression model appropriate for the outcome: a logistic model for binary disease status (e.g., disease vs. no disease), depicted as a sigmoid curve of disease status versus DNAmAge residuals; and a linear regression for continuous aging-related phenotypes, depicted as a scatter plot with a regression line. **Step 2** aggregates z-score estimates across multiple studies for the same phenotype using meta-analysis; the schematic shows a forest plot where each study’s z-score is combined to yield a pooled estimate. **Step 3** compares associations between all clocks and all phenotypes; a heatmap depicts the number of associations between epigenetic biomarkers and disease-related phenotypes, highlighting the breadth of the Associations analysis. The QR code directs to translage.io/assocations.

**Figure 9:**
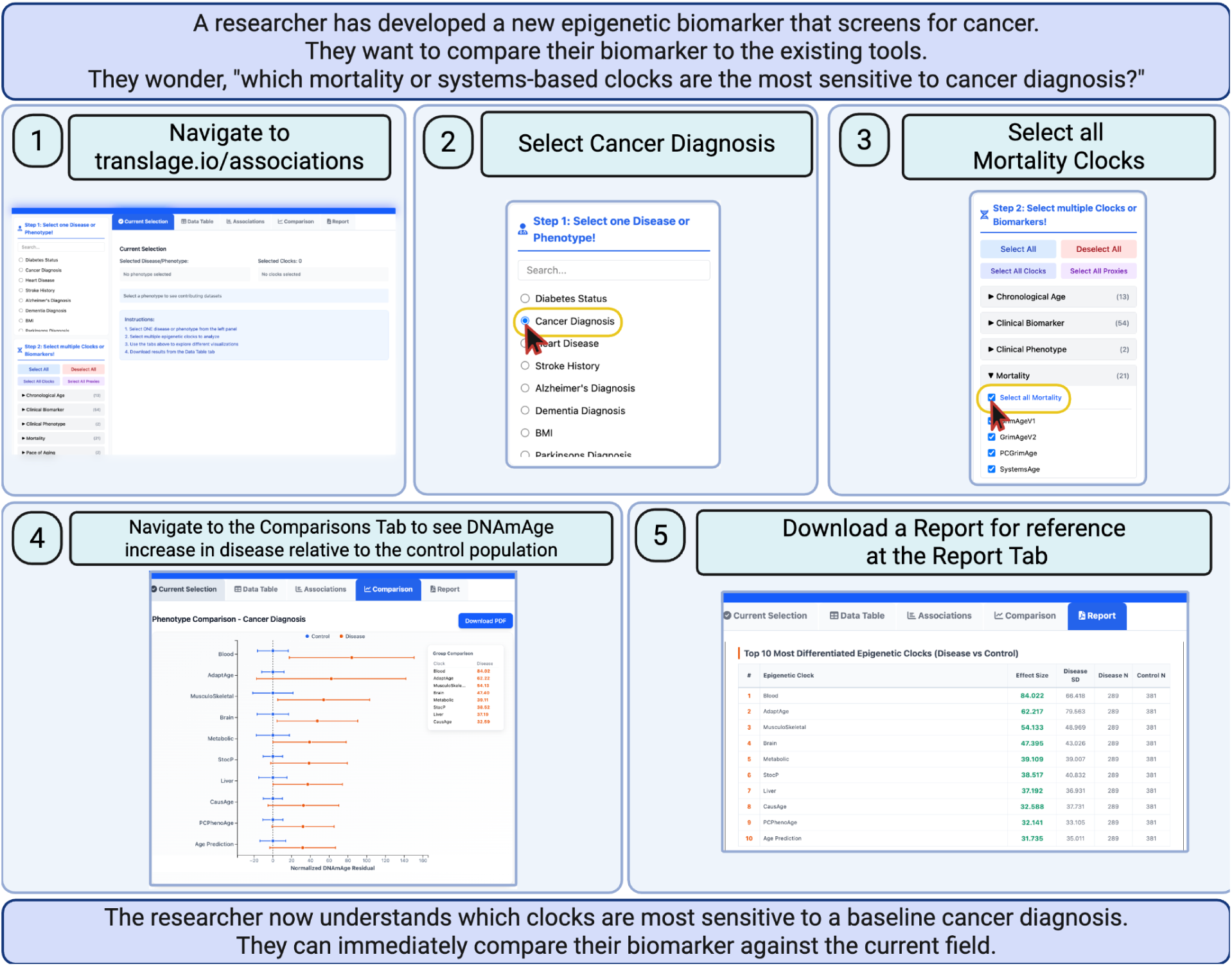
Associations Dashboard Use Case. A tutorial how-to for using the TranslAGE Associations Dashboard to find biomarkers most sensitive to baseline disease (in this case cancer). Screenshots are directly from the TranslAGE website. Yellow boxes and red-lined cursors are used to highlight important elements of the interactive page. Step 4 shows distributions of DNAmAge Residual in disease relative to the control population.

### Risk Prediction

Where the Association arm examines baseline relationships between epigenetic biomarkers and concurrent phenotypes, the Risk arm extends this framework to prognostic outcomes. In this analysis, we test whether baseline biomarker values predict future disease or mortality events using tools similar to the Associations analysis, but incorporating time-to-event models where applicable. A subset of datasets contained prospective phenotypic data for risk prediction analysis. Six datasets included future disease occurrence or mortality outcomes, with all six datasets sharing at least one overlapping prospective phenotype, enabling meta-analysis. In total, 81 dataset-phenotype pairs were identified for risk analysis, with seven future phenotypes measured in more than one dataset. Mortality outcomes (time-to-event with censoring) were available in one dataset – the Framingham Heart Study. Supplementary Table 3 contains details on prospective phenotype definitions. For future disease outcomes, we applied logistic regression (binary outcomes) or linear regression (continuous outcomes), adjusted for chronological age and sex. For mortality prediction, we employed Cox proportional hazards models to analyze time-to-event data, with DNAmAge residuals as predictors. Random-effects meta-analysis was performed to compute pooled effect sizes (log-odds ratios for disease, log-hazard ratios for mortality) across datasets measuring the same prospective outcome.

After meta-analysis, TranslAGE included 73 unique future disease phenotypes and mortality outcomes. Figure 10 summarizes the risk analysis workflow. These results can be explored through the Risk Dashboard at translage.io/risk, enabling users to query prospective disease prediction and mortality associations by phenotype or biomarker. Figure 11 showcases a use case of a researcher using the Risk Dashboard to identify a biomarker for predicting dementia onset.

**Figure 10:**
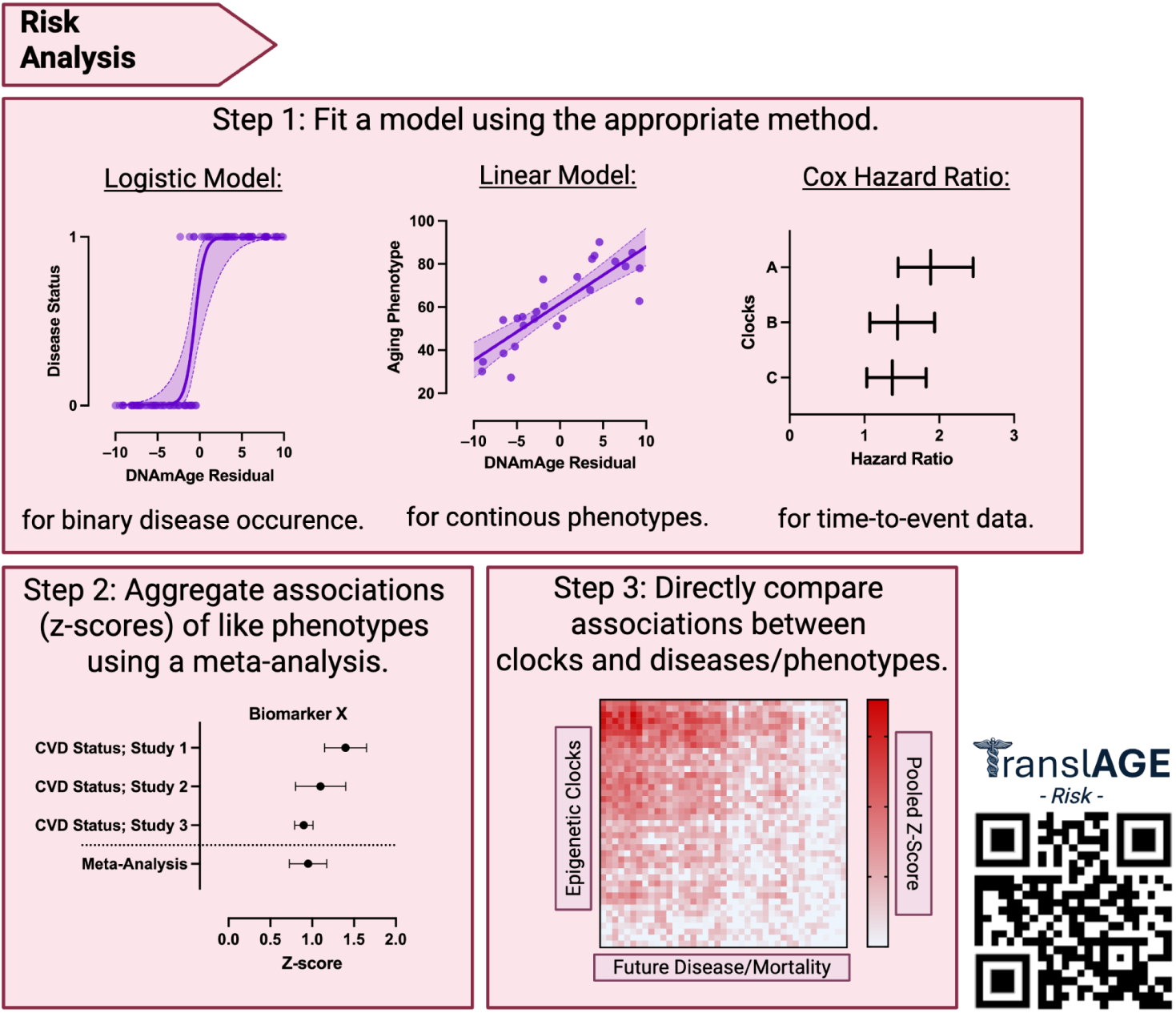
Risk Pipeline Methodology. This figure illustrates how TranslAGE evaluates biomarkers’ ability to predict future diseases or mortality. **Step 1** fits an appropriate model: a logistic model for binary outcomes (e.g., disease vs. no disease), a linear regression for continuous outcomes (e.g., future BMI), depicted as a scatter plot with a regression line, or a Cox Hazard Ratio with time-to-event information such as mortality. **Step 2** aggregates z-score estimates across multiple studies for the same phenotype using meta-analysis; the schematic shows a forest plot where each study’s z-score is combined to yield a pooled estimate. **Step 3** compares associations between all clocks and all future diseases or phenotypes. The QR code directs to translage.io/risk.

**Figure 11:**
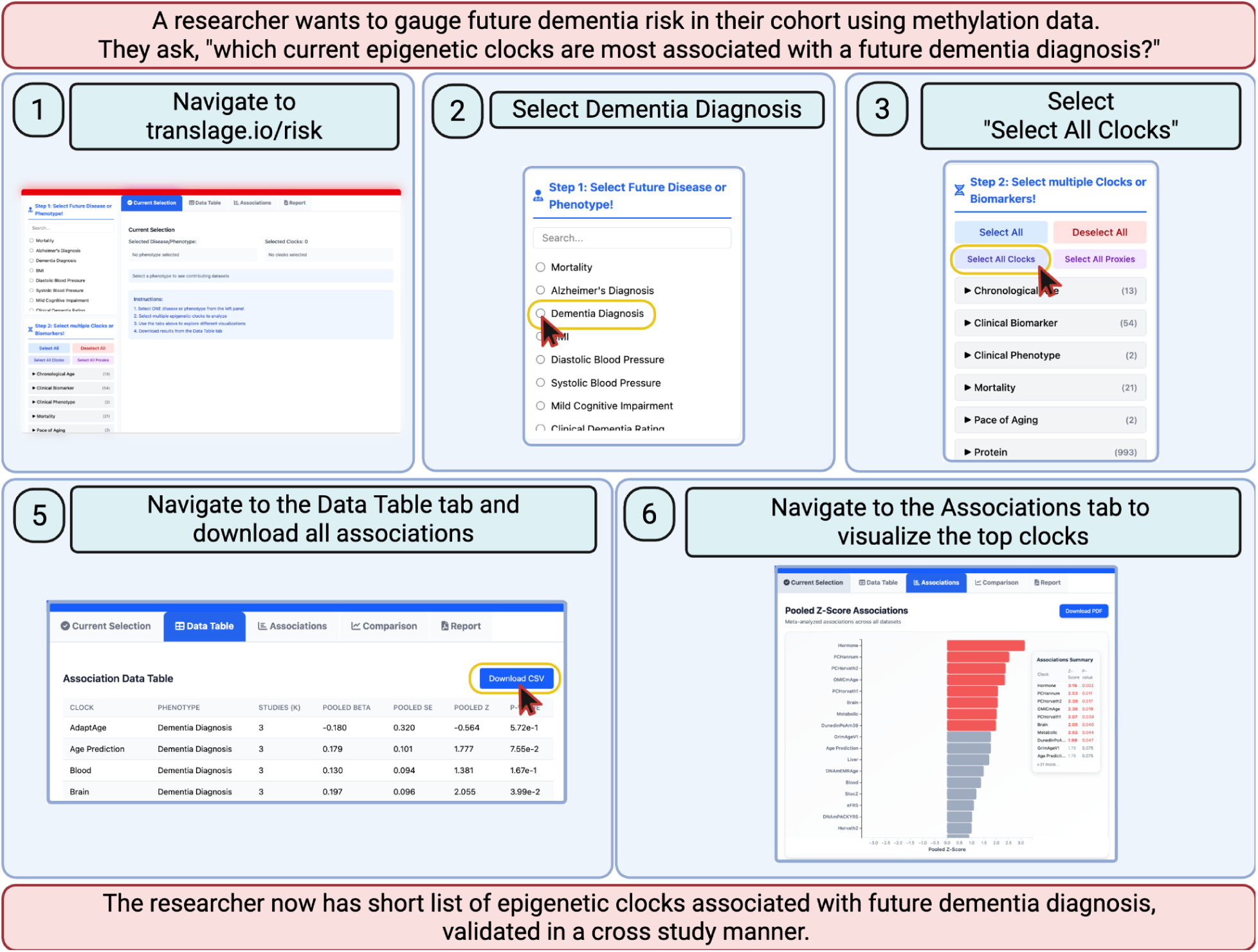
Risk Dashboard Use Case. A tutorial describing how to use the TranslAGE Risk Dashboard to select a biomarker for screening a cohort for future dementia risk using DNA methylation. Screenshots are directly from the TranslAGE website. Yellow boxes and red-lined cursors are used to highlight important elements of the interactive page.

### Explore Clocks with the Guiding STAR Dashboard

While the four domain-specific dashboards enable exploration of individual performance dimensions, real-world applications often require biomarkers that excel across multiple criteria simultaneously. To address this need, TranslAGE flexibly integrates the 4 STAR domains into a single composite score for any biomarker, tailored to user priorities, and provides an interactive dashboard for exploring the results (translage.io/recommend).

The tool allows users to specify which STAR domains are relevant to their study design and returns a ranked list of biomarkers optimized for those criteria. Users can filter domains further for context-specific biomarker selection. Association can be filtered for disease category (Cardiovascular, Metabolic, Neurological & Mental Health, Cancer), Treatment response can be filtered for event subtype (Lifestyle, Pharmacological, Supplement, or Event), and Risk can be filtered for disease or mortality only. Interactive score breakdowns provide full transparency into how each biomarker’s composite score was calculated.

We calculated a single composite STAR score for each biomarker by combining four domain-specific z-scores. The Stability domain derived z-score from the pooled ICC estimates, Treatment response uses scaled effect size, and Associations and Risk both use the pooled effect size and standard errors. For further details on computing a z-score for each domain, see *Methods*.

To demonstrate the utility of multi-domain STAR integration, Figure 12 showcases how a researcher planning a clinical trial can harness all the domains of TranslAGE to select the best epigenetic biomarker for their needs. In this case, the researcher selects all 4 domains of TranslAGE, and therefore higher STAR scores indicate a greater reliability, responsiveness, disease association, and predictive utility.

**Figure 12:**
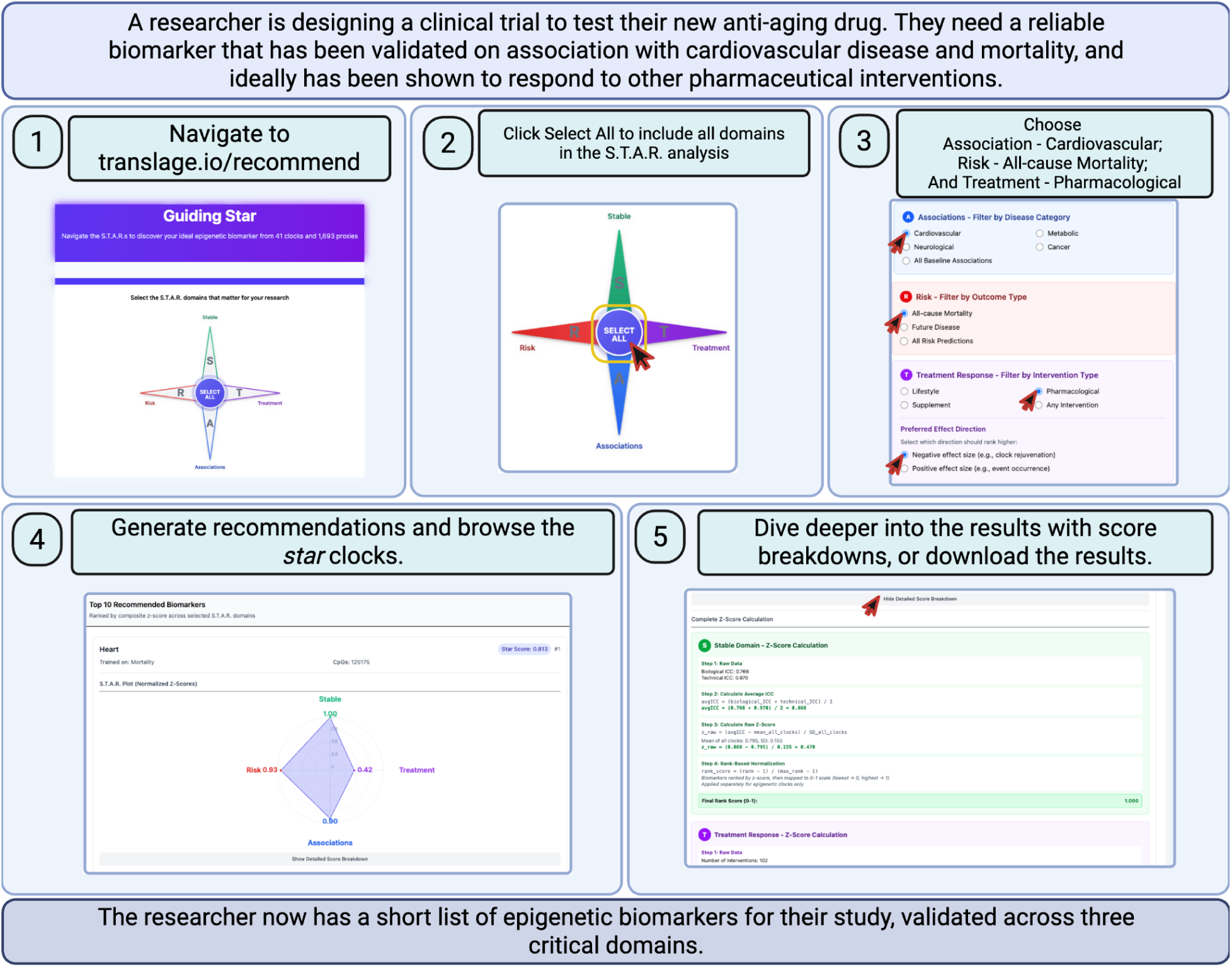
Guiding STAR Dashboard Use Case. Tutorial on how to use the Biomarker Recommendation Wizard. Screenshots are directly from the TranslAGE website. Yellow boxes and red-lined cursors are used to highlight important elements of the interactive page.

### Scaling TranslAGE

TranslAGE was designed to scale along two key dimensions: the number of datasets it integrates and the number of clocks it computes.

New datasets can be incorporated after they have been cleaned and harmonized. Once screened, the datasets can be flagged and then routed to one or more of the four STAR domains according to the predefined criteria. The resulting outputs can be published directly to the TranslAGE dashboards, making new findings available to the community immediately. This streamlined integration not only expands the resource but also benefits collaborators by returning 41 clock scores and benchmarking their results against comparable studies within TranslAGE.

New clocks can also be incorporated into TranslAGE for immediate, independent validation. This enables side-by-side comparison of novel biomarkers with established clocks while simultaneously generating new metrics across all datasets already housed in TranslAGE. To pilot this process, we collaborated with Carreras-Gallo et al. (2024), who developed a large compendium of 1,694 epigenetic biomarker proxies (EBPs) predicting proteins, metabolites, and other clinical biomarkers from DNA methylation data (Carreras-Gallo et al. 2024). Incorporating these EBPs into TranslAGE allows for their independent validation and comparison against existing clock models. All EBPs are fully integrated into the TranslAGE dashboards and can be explored at any time by filtering for Proxies rather than Clocks.

## Discussion

TranslAGE represents a foundational step toward a standardized ecosystem for validating biological aging biomarkers. By harmonizing 179 methylation datasets, and evaluating clocks across the four STAR performance domains, the platform provides an unprecedented scope of biomarker validation across diverse populations and experimental contexts.

TranslAGE provides unique value to the geroscience and aging biomarker community in three ways. First, it offers a rich collection of harmonized methylation datasets for exploration and rapid download. Second, it delivers a comprehensive suite of precalculated aging biomarker scores that can be explored interactively or downloaded for further analysis. Third, it demonstrates a systematic analysis that highlights the strengths and limitations of the existing epigenetic clocks. This allows users to select the best clock for their particular use case, and also lays the groundwork for continual improvement of clocks. Together, these three facets provide a massive foundation for advancing the study, development, and application of epigenetic aging biomarkers.

Several existing platforms have advanced the computation, visualization, and benchmarking of epigenetic clocks, but these mostly remain limited to clock calculation and associations with phenotypes. TranslAGE advances beyond these efforts by harmonizing 179 methylation datasets and evaluating 41 clocks and 1,694 proxies within its S.T.A.R. framework to enable systematic, cross-study benchmarking of epigenetic aging biomarkers in diverse contexts. The particular strength is the inclusion of 129 longitudinal datasets that allow for calculation of Stability and Treatment response metrics in parallel to the more common Association and Risk prediction metrics. This scope and design establish TranslAGE as the most comprehensive resource to date for evaluating the performance and translational potential of epigenetic clocks.

Harmonization is essential because methylation datasets originate from multiple platforms and studies with heterogeneous metadata. TranslAGE’s standardized columns, residualization process and unified statistical frameworks ensure that associations and effect sizes are comparable across datasets. The interactive dashboards, which allow users to filter by their relevant criteria and requirements, make exploratory analyses practical for non-bioinformaticians. This interactive approach accelerates hypothesis generation and validation, a critical need in aging research where subtle effects and confounders can otherwise be overlooked. By offering precomputed clock scores and large-scale comparisons, TranslAGE lowers the barrier to testing existing and emerging epigenetic clocks. Researchers can quickly identify clocks for their specific needs, including diagnostics, clinical trial planning, or disease screening.

One critical caveat of TranslAGE is the lack of multiple-hypothesis testing implementation in its current release. TranslAGE involves hundreds of thousands of statistical tests and uses no multiple-testing correction. Therefore, individual findings pulled out of the context of the broader discoveries within this resource need to be investigated further before any conclusions can be drawn. Indeed, TranslAGE was in part inspired by the goal of drawing attention to the risk of selection bias with epigenetic clocks. TranslAGE is a resource for looking at trends, and for discovering (or questioning) the potential of epigenetic clocks as aging biomarkers. For these purposes, multiple-hypothesis testing is not necessary at this current stage. However, it is our plan to include a feature at a later date to rapidly switch the statistical analysis to include different methods of multiple-hypothesis testing (Benjamini-Hochberg, Bonferonni) for users to explore. It is important to note that the very purpose of TranslAGE is to help researchers identify which clocks are most appropriate for their particular use case, based on prior data and STAR metrics. By limiting analysis to a small panel of the most appropriate clocks, researchers can minimize multiple testing in their own studies.

Additional limitations in our approach include the issue of batch effects, including the cross comparison between studies and arrays without explicit batch correction. However, it should be noted it is our explicit intention to develop STAR scores on both 450K and EPIC arrays, to systematically understand how algorithms perform on different arrays. Nevertheless, there are likely some benefits to correcting for array (Fortin, Triche, and Hansen 2017; Lussier et al. 2024; Tay et al. 2025; Zhuang et al. 2025), and users should be aware that array may influence STAR metrics and interpret the results accordingly based on the annotated array type. Additional preprocessing steps in the initial methylation array fluorescence also differed by study, and were not entirely accounted for. These differential preprocessing steps include probe filtering, the choice of background correction technique (NOOB or ENmix), dye bias correction. This may be addressed by directly comparing results from a dataset already preprocessed by the original author compared to a dataset preprocessed by a standardized pipeline, and determining to what extent STAR metrics are affected by preprocessing.

Another limitation in the current TranslAGE framework is the heterogeneity in phenotype reporting. We used a manual classification schema based on dataset documentation (Supplementary Table 3), however there is an unavoidable level of incongruence between the diagnostic routine of different datasets. Phenotype harmonization will be particularly important as TranslAGE expands the Association and Risk branches with additional large cohort studies.

Notably, we deliberately limited TranslAGE’s initial release to human blood-based DNA methylation due to the availability of harmonizable longitudinal data, which maximizes TranslAGE’s utility for clinical translation. Blood is critical for aging biomarkers because blood can be repeatedly sampled and assessed for stability over time and responsiveness to interventions. Furthermore, blood can be banked and assayed in longitudinal aging cohorts with follow-up assessments for aging outcomes many years later. Without such follow-up data, future risk prediction is not possible. While chronological age predictors can be trained using cross-sectional data, these suffer many drawbacks including statistical limitations, period effects, and false positive changes in intervention studies (Sluiskes et al. 2024; Nelson, Promislow, and Masel 2020; Schaie 1967; Borrus et al. 2024; Sehgal et al. 2024). However, there are many other applications for TranslAGE beyond clinical translation. As TranslAGE is indefinitely extensible, the broader vision is to expand into additional tissues, species, and biomarker types (including clinical, transcriptomic, proteomic, and metabolomic algorithms and datasets).

Additional avenues for expansion lie in automation and real-time integration. The current pipeline harmonizes datasets manually, but automated ingestion of newly published methylation studies would accelerate growth and ensure continual updating of the resource. Implementing automatic scraping, workflows, and quality control pipelines could link TranslAGE to public repositories such as GEO, allowing the resource to serve as a live, continuously updated benchmarking environment.

Ultimately, TranslAGE aims to become a shared infrastructure for the aging-biomarker community—supporting reproducible, large-scale validation and unifying disparate datasets under a single analytic framework. By providing harmonized data, transparent pipelines, and interactive exploration tools, TranslAGE bridges discovery and translation, enabling researchers and clinicians alike to identify STAR biomarkers of biological aging. As TranslAGE expands to include additional data and statistical models, the platform will continue to evolve toward a comprehensive living atlas of aging biology.

## Methods

### Dataset curation and harmonization

Methylation datasets were screened to ensure they contained realistic beta-value matrices or raw .idat files derived from human blood samples measured on Illumina methylation arrays. Datasets required a minimum of 10 samples and at least 95% probe coverage. Datasets were harmonized by hand by our team. Curated information was derived from available phenotypic columns, and formatted to fit into TranslAGE structure and variable types (i.e., characters, floats, factors). All methylation datasets were mean imputed, such that missing values for a given individual were imputed from the datasets at that CpG site. Entirely missing probes were replaced with 0s. A list of all datasets included in TranslAGE at the time of submission see Supplemental Table 1.

### Clock calculations

We utilized the methylCIPHER package (https://github.com/HigginsChenLab/methylCIPHER) (Thrush et al. 2022) and, following the package’s structure, added additional clocks and epigenetic-based biomarker algorithms to fit our needs. For each dataset, each clock received an identical methylation array to avoid discrepancies that arose from different algorithm inputs. A list of all clocks and biomarkers used in this analysis at time of submission can be found in Supplemental Table 2.

Three datasets out of 179 were missing chronological age. In these fringe cases, we made note of the missing information, and substituted in Zhang2019 (Zhang et al. 2019) for the missing age to complete our analysis. To avoid confounding our findings, Zhang2019 is omitted from the clock validation analysis pipelines. No explicit batch correction across arrays was performed.

### Age residual

DNA methylation clock residuals were calculated by regressing each epigenetic biomarker estimate against chronological age to remove age-related variation. For datasets lacking chronological age data (3 datasets total), the Zhang2019 epigenetic age estimate was used as a proxy for age in the residualization models.

### Association and Risk Pipelines

Datasets were screened for phenotypic columns related to aging health and disease. Datasets containing relevant information were flagged for the risk or association analysis. Supplementary Table 3 has the definitions used for categorizing and combining phenotypic/disease columns across datasets.

For each epigenetic clock-phenotype combination, appropriate regression models were automatically selected based on outcome type. Binary disease outcomes were analyzed using logistic regression (’glm(…, family=binomial)’), continuous phenotypes using linear regression (’lm(…)’), ordinal outcomes using proportional odds models (’polr(…)’), and survival outcomes using Cox proportional hazards models (’coxph(…)’). All models included standardized clock values (’scale(clock)’) and were adjusted for chronological age and sex. From each fitted model, we extracted the regression coefficient (β), standard error (SE = |β/z|), and test statistic (z-score). For clock-phenotype combinations with multiple studies, random-effects meta-analysis was performed using the ‘metafor’ package (’rma(yi = beta, sei = se, method = “REML”)’), implementing the model β_i_ = θ + u_i_ + ε_i_, where β_i_ is the effect size from study i, θ is the pooled effect size, u_i_ ∼ N(0, τ²) represents between-study heterogeneity, and ε_i_ ∼ N(0, SE_i_²) represents within-study sampling error.

### Treatment Response Analysis Pipeline

Datasets containing longitudinal samples were flagged for Treatment response analysis if A) the datasets had more than 5 subjects (10 samples) and B) the follow up samples were taken more than 48 hours after the baseline samples. Intervention responsiveness was assessed using paired t-tests comparing pre- and post-intervention epigenetic clock values within individuals. Effect sizes were scaled relative to a large EPICv1 study to enable cross-study comparisons. For each clock-intervention combination, we calculated the mean change in clock residuals and transformed this to a standardized effect size using the reference dataset biomarker age-adjusted standard deviation as the reference scale.

### Stability Analysis Pipeline

Datasets with replicates were flagged for stability analysis. Samples taken from the same individual with the same blood draw were tagged as technical replicates. Conversely, samples from an individual within a 24 hour window were marked as biological replicates. Clock reliability was assessed using Intraclass Correlation Coefficients (ICCs) calculated from mixed-effects models (’lmer(y ∼ 1 + (1|SUBJ))’), where Y_i_□ = μ + u_i_ + ε_i_□, with u_i_ ∼ N(0, σ_u_²) representing between-subject variation and ε_i_□ ∼ N(0, σₑ²) representing measurement error. ICC was computed as σ_u_²/(σ_u_² + σₑ²). To combine ICC estimates across datasets, we implemented random-effects meta-analysis using Fisher’s z-transformation for approximate normality (’z = ½ ln((1 + ICC)/(1 -ICC))’), with variance ‘Var(z) = 1/(n-3)’. The meta-analysis model z_i_ = θ + u_i_ + ε_i_ was fitted using ‘rma(…, method = “REML”)’, where θ represents the overall effect size, u_i_ ∼ N(0, τ²) represents between-study heterogeneity, and ε_i_ ∼ N(0, v_i_) represents within-study variance. Pooled ICC estimates were back-transformed using ‘(exp(2θ^) - 1)/(exp(2θ^) + 1)’.

### Unified STAR Dashboard

Domain-specific z-scores for the composite STAR score (Stability, Treatment response, Associations, Risk) for each biomarker were calculated as follows. For the Stability domain, an average ICC (*avgICC)* was calculated as the average of biological and technical ICC. Then, z-scores were derived as

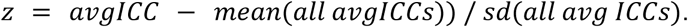

For the Treatment response domain, filters for intervention type were applied if specified. Then for each clock, a z-score was calculated using scaled estimates, where

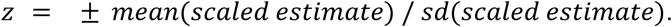

The calculation can be negated if the user is more interested in decreasing effect sizes (decreasing biological age; a beneficial response) with a toggle option on the dashboard. For the Associations and Risk domain, z-scores were calculated from pooled effect size (β) and standard error *se*, where

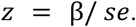

z-scores were filtered by any user-selected disease category, and then aggregated using Stouffer’s method for combining z-scores across studies,

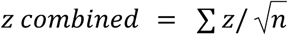

where *n* is the number of z-scores being aggregated. For each domain, rank-based normalization maps z-scores to a 0 to 1 scale, where biomarkers are ranked by performance and evenly distributed (lowest z-score = 0; highest = 1), ensuring all domains contribute equally to the final score.

The composite STAR score aggregates the z-scores from the selected domains using a simple average, to ensure equal contribution of the four domains. Only domains selected by the user contributed to the final STAR score. All clocks and biomarkers were ranked by STAR score, and results displayed a breakdown of the contributing domains.

### Data and Code Accessibility

All analyses were performed in R (v4.3.2; packages: methylCIPHER v1.2.1, metafor v4.4-0, lme4 v1.1-35, survival v3.5-7). Analysis datasets are available upon request. Cleaned datasets and related calculated clock scores can be downloaded directly from the TranslAGE website, in the Front Page Dashboard (pending publication), if the original datasets are also publicly available. Private or protected datasets have variable levels of public accessibility depending on data use agreements on a dataset-by-dataset basis. Limitations on data accessibility is reported on a per-dataset basis after a data download request is made on the Front Page Dashboard. To protect the work, full dataset download functionality is limited until this manuscript is accepted in a peer reviewed journal. All non-restricted data and derived results will be accessible through TranslAGE upon acceptance. Website screenshots taken October 2025. Figures for this publication were created using BioRender.com.

## Acknowledgements

This work is supported by the National Institute on Aging (NIA:1R01AG065403 to A.H.C.). DSB is supported on a T32 NIA training grant (NIA:T32AG019134). The work is also supported by the Impetus Grant (R.S.), the Gruber Science Fellowship at Yale University (R.S., J.F.A), and the Thomas P. Detre Fellowship Award in Translational Neuroscience Research from Yale University (to A.H.C.). The authors would like to thank the researchers and clinicians who made their study data publicly available, as without their data this project would not have been possible. All figures were created in BioRender.com.

## Author Contributions

D.S.B., R.S., and A.H.C conceived the project and study design. D.S.B., R.S., J.F.A., J.G., G.Z., J.K., Y.M., and A.H.C. contributed to data curation and harmonization. D.S.B., R.S., and J.K. expanded MethylCIPHER to include additional algorithms. J.L.S. provided additional epigenetic biomarker proxies and planning for TranslAGE expansions. D.S.B. built the pipeline backend and website. D.S.B. and A.H.C. wrote the manuscript, and all authors reviewed and contributed to the manuscript.

